# Transgenic resources for functional studies on the *hsrω* gene in *Drosophila*

**DOI:** 10.1101/2021.12.26.474178

**Authors:** Ranjan Kumar Sahu, Deepti Trivedi, Subhash Chandra Lakhotia

## Abstract

The *hsrω* gene of *Drosophila melanogaster*, an early discovered non-coding developmentally active and stress-inducible gene, has pleiotropic actions through its multiple nuclear and cytoplasmic non-coding transcripts. With a view to understand this diverse functions and mechanisms of actions of this gene, we generated a series of transgenic lines for over-expression or RNAi of different transcripts and CRISPR-Cas9 mediated deletion alleles. These are described here.

## Introduction

The *hsrω* gene, located at the 93D cytogenetic region in *Drosophila melanogaster*, is an early discovered long non-coding RNA gene which is developmentally active, stress responsive and expressed in almost all cell types (Lakhotia, 2011). Its non-coding nature and singular inducibility by diverse amides (Lakhotia and Mukherjee, 1970; Tapadia and Lakhotia, 1997) makes it different from other stress responsive genes. All species of *Drosophila* carry an equivalent of *hsrω*, with remarkable conservation of its architecture and inducibility amidst high sequence diversification (Sahu *et al*., 2020).

The *hsrω* gene’s conserved architectural features (Fig. 1a) include unique sequence encompassing two exons and a more conserved omega intron in its proximal part whilethe distal region typically includes tandem repeat sequences (Sahu *et al*., 2020). A conserved short ORF (ORFω), with a potential to code for a short peptide is present within the exon 1in different species (Sahu *et al*., 2020).

**Fig. 1.**
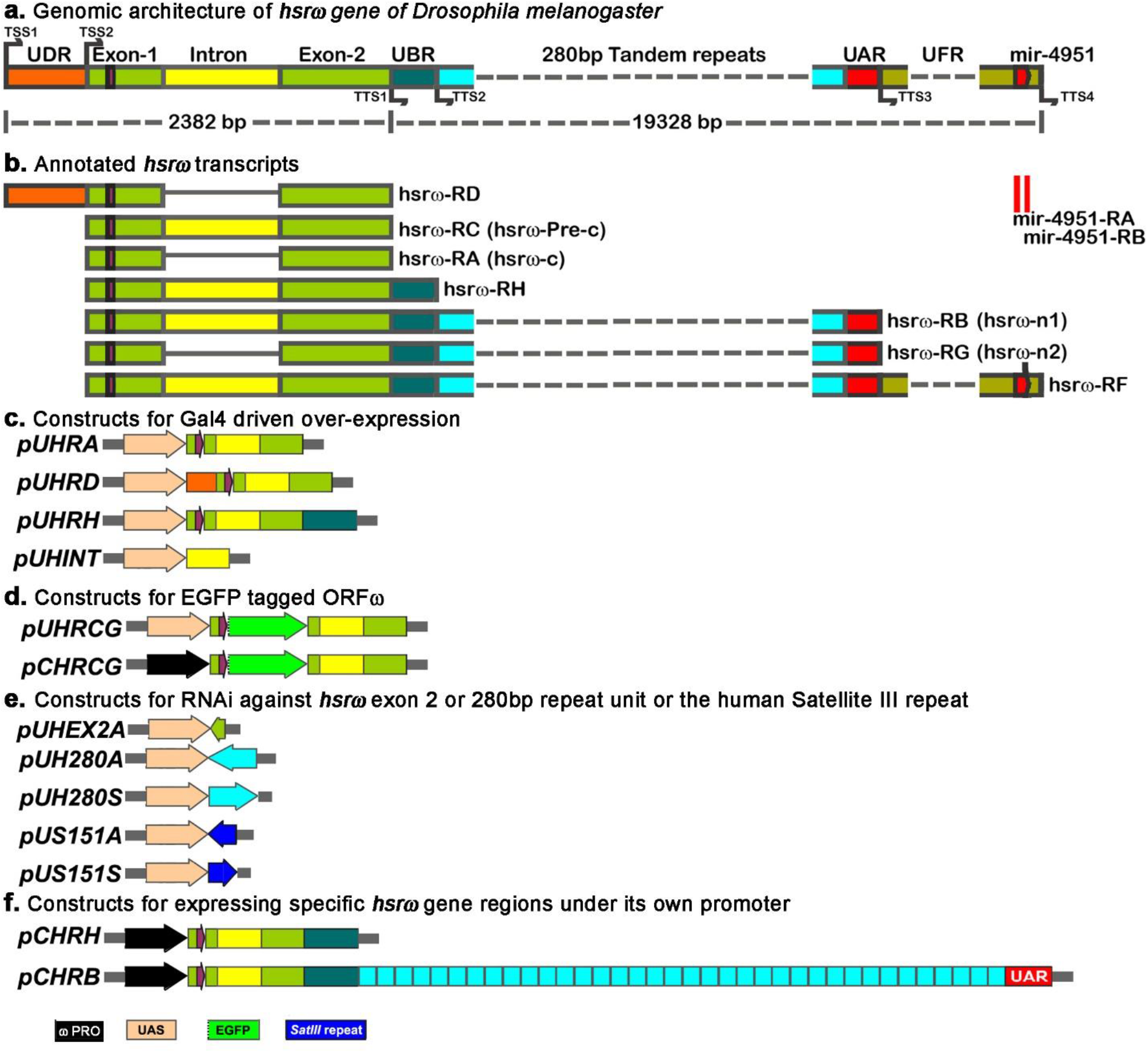
Genomic coordinates of *hsrω* gene in *D. melanogaster* and the prepared transgenic constructs. **a.** Organization of the *hsrω* gene (adapted from www.flybase.org): two upward arrows indicate the two transcription start sites (TSS1 and TSS2) while the four downward arrows indicate the four annotated transcription termination sites (TTS1–TTS4); different segments of the gene are color-coded and named (noted above the segment) following (Sahu *et al*., 2020). **b.** Transcripts produced by the *hsrω* gene: the spliced out intron (yellow) is indicated by a black line joining the exons 1 and 2 regions (green boxes); the transcripts are named (on right of each transcript) following www.flybase.org (transcript names in parentheses are the common names in published literature), **c.** Constructs for expression of different regions of *hsrω* downstream of the UAS promoter (in *pUAST* vector). **d.** Constructs for expression of ORFω-EGFP fusion protein under the UAS promoter (in *pUAST* vector) or 4.3kb *hsrω* promoter (in *pCaSpeR5* vector). **e.** Constructs for inducing RNAi against exon 2 or repeat unit of *hsrω* gene or against the human Sat III transcripts in *pUAST* vector. **f.** Constructs in *pCaSpeR* vector carrying 4.3 kb (in *pCHRH* clone) or ~5kb (in *pCHRB* clone) of *hsrω* promoter (which also includes the UDR region of the RD transcript) and the indicated downstream transcribed sequence. Additional color codes for the *hsrω* promoter (*ω* PRO), *UAS* promoter (in the *pUAST* vector), the EGFP coding region (without its start AUG codon) and the human *SatIII* repeat are indicated at bottom of the figure.

The Flybase annotation indicates that the *hsrω* gene in *D. melanogaster* produces a total of seven transcripts of which some are nuclear while at least one of them is known to be cytoplasmic. The distal region includes a long stretch of 280bp tandem repeats. A conserved miRNA gene (*mir-4951*) is present at the 3’-end of the *hsrω* gene (Sahu *et al*., 2020). This gene’s nuclear transcripts carrying the distal tandem repeats organize the nucleoplasmic omega speckles that regulate dynamics of hnRNPs and some other RNA binding proteins in cells under normal and stressed conditions (Prasanth *et al*., 2000; Singh and Lakhotia, 2015).

With a view to understand the diverse functions of this gene’s multiple transcripts, we generated several transgenic *D. melanogaster* lines, using P-element based vectors. Two types of P-element vectors viz., *pUAST*(Giordano *et al*., 2002) and *pCaSpeR5* (Thummel *et al*., 1988) were used. The *pUAST* vector was used for targeted expression of the sequence of interest with a GAL4 driver while the *pCaSpeR5* was used for expression of the specified *hsrω* gene sequence under its own promoter. In addition, two CRISPR/Cas9 mediated deletion alleles have also been generated.

## Methods and Results

### A. Constructs to conditionally express different transcripts of the *hsrω* gene

In order to understand functions of three diverse transcripts ranging in size from ~1.2kb to ~21.2kb (Fig. 1b), different *hsrω* transcript producing regions were placed under the UAS promoter in *pUAST* vector.

i. ***pUHRA***: Due to unavailability of PstI in MCS of the *pUAST* vector, the 1.2kb PstI digested fragment from the *pJG10* plasmid (from M. L. Pardue’s lab carrying the 1.2 Kb RA cDNA in *pSP65* vector) was initially inserted in *pBlueScript SK(-)* in forward orientation and named *pBSHRA*. The EcoRI + XbaI fragment from *pBSHRA* was ligated in the *pUAST* vector at EcoRI and XbaI sites to generate the *pUHRA* construct (Fig. 1c).
ii. ***pUHRD:*** A DNA fragment from TSS1 to TTS1 (~2.4kb) along with the intron segment was PCR amplified using the primers RDF and 2R (Table 2) with EcoRI and XbaI overhangs, respectively. After double digestion with EcoRI and XbaI, the amplified fragment was ligated in the *pUAST* vector to generate the *pUHRD* construct (Fig. 1c).
iii. ***pUHRH***: The DNA fragment from TSS2 to TTS2 (~2.8kb) was amplified through PCR using the primers pairs 1F and RHR (Table 2) with EcoRI and XbaI overhangs, respectively. Following double digestion of the amplified fragment with the two restriction endonucleases, it was ligated with EcoRI and XbaI digested *pUAST* to obtain the *pUHRH* construct (Fig. 1c).
iv. ***pUHINT***: For this construct, the *hsrω* intron (712bp) was amplified using the primers INTF and INTR (Table 2) with EcoRI + XbaI flanking sites. The resulting amplicon was double digested with the two enzymes and ligated with EcoRI and XbaI digested *pUAST* to produce the *pUHINT* construct (Fig. 1c).

### B. Transgenic constructs to detect the translatability of ORFω

i. ***pUHRCG***: In order to detect translatability of the short ORFω in exon 1, EGFP reporter sequence was inserted in frame downstream of the ORFω after removing the stop codon of ORFω and initiation codon of the reporter gene. To achieve this three PCR amplified DNA fragments were ligated. Two fragments were amplified from the *hsrω* gene and the third (EGFP fragment) from the *pEGFP-N1* plasmid (Clontech; Kang *et al*., 2001). The first *hsrω* gene fragment was amplified from TSS2 to 3’ end of ORFω (excluding its stop codon) with primers 1F and 1R (Table 2) with EcoRI and SpeI overhang sites, respectively. The second fragment downstream of ORFω to TTS1 was amplified with primers 2F and 2R (Table 2) with KpnI and XbaI overhang sites, respectively. The EGFP CDS (excluding its initiation codon) was amplified from the *pEGFP-N1* plasmid with primers EGFPF and EGFPR (Table 2) having SpeI and KpnI overhang sites, respectively. Finally, the three amplicons were digested with respective sets of enzymes and ligated to form a linear EGFP tagged ORFω as part of the *hsrω-RC* transcript which was finally ligated into *pUAST* vector at EcoRI and XbaI sites to generate the *pUHRCG* plasmid (Fig. 1d). Appropriate in-frame fusion of the ORFω with EGFP in the *pUCHRG* construct was confirmed by dideoxy sequencing using the sequencing primers pUASTF and pUASTR, (Table 3) corresponding to the vector backbone.
ii. ***pCHRCG:*** The EcoRI + XbaI fragment used for generating the above *pUHRCG* plasmid was also cloned in *pCaSpeR5* and screened for positive clones. The 4.3kb region upstream of the TSS2 of *hsrω* gene was amplified with the primers HPF and HPR (Table 2) having EcoRI sites at both the ends. The EcoRI digested 4.3kb promoter sequence carrying amplicon was inserted into the *pCaSpeR5+HRCG* plasmid at EcoRI site and the resulting plasmid (*pCHRG*) was used to transform *DH5α*. To screen clones (*pCHRCG*) with correct orientation (Fig. 1d) of the promoter, plasmids from the transformed colonies were digested with XbaI and selected for those showing restriction pattern of two bands of 7931bp and 6992bp sizes, respectively. The correct orientation was further confirmed by di-deoxy sequencing of the insert and ligation junction using sequencing primers pC5F and pC5R (Table 3) corresponding to the vector backbone.

### C. Transgenic constructs for over-expression of sense and antisense transcripts corresponding to the 280bp tandem repeat unit of *hsrω* and 158bp repeat unit of human *SatIII* transcripts

The widely used RNAi line against the nuclear transcripts of *hsrω*, generated earlier in our lab using the 280bp repeat cloned in the *pSympUAST-w* vector (Mallik and Lakhotia, 2009), produces sense and anti-sense transcripts under the UAS promoter and down-regulates the *hsrω* nuclear transcripts. We now generated two new RNAi lines in *pUAST* vector that can be used for down-regulation of cytoplasmic and nuclear transcripts, respectively, and one line that over-expresses the sense strand of the 280bp repeat unit. In addition, we also generated two transgenic lines that express the sense or anti-sense strand of the human *SatIII* repeats, which are considered to be functional homologs of the *Drosophila hsrω* nuclear transcripts (Jolly and Lakhotia, 2006), under the UAS promoter.

i. ***pUHEX2A***: Complete exon-2 region was aligned against transcript database of *Drosophila melanogaster* using the BLAST tool. A unique 80bp fragment from 3’ end of Exon-2 was selected and PCR amplified using wild type genomic DNA and the primers EX2AF and EX2AR (Table 2) having EcoRI and XbaI overhangs, respectively. The PCR fragment was digested with respective restriction endonucleases and inserted into the *pUAST* vector to produce the *pUHEX2A* construct (Fig. 1e)
ii. ***pUH280A***: In order to clone the 280bp repeat of *hsrω* in antisense orientation in *pUAST* vector, the *pDRM30* plasmid clone (from M. L. Pardue’s Lab), carrying 280bp Asu II fragment of *hsrω* repeat in *pGEM3* vector was double digested with PstI + XbaI to release the 280bp fragment which was re-cloned into *pBlueScript SK(-)* and named as *pBS280*. The NotI + XhoI fragment from *pBS280* was ligated into the NotI + XhoI digested *pUAST* vector to generate the *pUH280A* construct (Fig. 1e).
iii. ***pUH280S***: The above *pBS280* was double digested with EcoRI + XbaI to release the 280bp repeat fragment which was re-cloned in the *pUAST* vector to generate the *pUHS280S* construct (Fig. 1e)
iv. ***pUS158A***: In order to clone the 158bp repeat of *SatIII* in antisense orientation in *pUAST* vector, the ~160bp EcoRI + BamHI digested fragment from *pGEM2-98* clone, harboring a 158-bp fragment of the human *SatIII* repeat from the 9q12 locus in *pGEM2* vector (a kind gift from Dr. Caroline Jolly, INSERM, France) was inserted into the *pBlueScript SK (-)* vector, and named as *pBS158*. The NotI + XhoI fragment from *pBS158* was subcloned in the *pUAST* vector to produce the *pUS158A* construct (Fig. 1e) that can down-regulate the human *SatIII* transcripts.
v. ***pUS158S***: In order to generate this clone, the EcoRI + XbaI digested fragment from the above *pBS158* was re-cloned in *pUAST* at EcoRI + XbaI sites to generate the *pUS158A* transgene construct (Fig. 1e), for overexpression of the human *SatIII* repeat.

### D. Transgenic constructs to generate *hsrω*-RH and RB transcripts under its own promoter

i. ***pCHRH:*** In order to generate this clone, the EcoRI + XbaI digested ~2.8kb fragment from the above described *pUHRH* plasmid was ligated into EcoRI + XbaI digested *pCaSpeR5* and screened for the positive clones (*pCaSpeR5+HRH* clone). The 4.3kb promoter region of the *hsrω* gene (as described above for the *pCHRCG* clone) was ligated at the EcoRI site of the *pCaSpeR5+HRH* clone to generate the *pCHRH* plasmid (Fig. 1f). To identify the clones with right orientation of the promoter with respect to the HRH sequence, isolated *pCHRH* plasmid DNAs from different transformed colonies were digested with XbaI to identify colonies that carry plasmid which generates two fragments of 7040bp and 7931bp, respectively.
ii. ***pCHRB:*** To generate this construct, the *pDRM102* plasmid (from M. L. Pardue’s lab) was digested with EcoRI + BamHI to generate three fragments of ~19kb, ~1kb and ~2.2kb sizes. The ~19kb fragment (corresponding to the *hsrω-RB* transcribing region and ~5kb upstream endogenous promoter region) was purified, ligated into the *pCaSpeR5* at EcoRI and BamHI sites to generate the *pCHRB* construct (Fig. 1f) and transformed into DH5α. For confirming positive colonies, the isolated plasmids were double digested with PstI + BsaI to get five fragments of 1935bp, 2229bp, 3445bp, 6612bp and 12768bp, respectively.

The above plasmid constructs were used to transform DH5α and the purified plasmid DNAs in each case were confirmed to carry the desired transgene by restriction digestion and/or dideoxy sequencing. The purified DNAs were used for microinjection in *w*^-^ embryos together with the helper p-transposase-encoding plasmid DNA at the Fly-facility of the Bangalore Life Science Cluster, Bangalore. The emerging flies were crossed with *w*^-^ flies. The progeny flies with red-eyes (due to germline transformation with the transgenic plasmid carrying the *mini-white* reporter) were selected and crossed with *w-; Sp/CyO; TM3/TM6B* double balancer stock to determine the chromosomal insertion of the transgene and subsequent establishment of transgenic stocks. In some cases, the transgene insertion was associated with recessive lethality (see Table 1). In such cases, the transgene carrying parent was crossed with *w*^-^; +/+; +/+ flies and the red-eyed progenies were crossed with white-eyed (*w*^-^; +/+; +/+) flies for three generations to remove potential 2^nd^ site mutation that may be responsible for the recessive lethality. In several cases, free-floating for three generations produced transgene-carrying homozygous viable lines, which are maintained as stocks for further studies (Table 1).

**Table 1.**
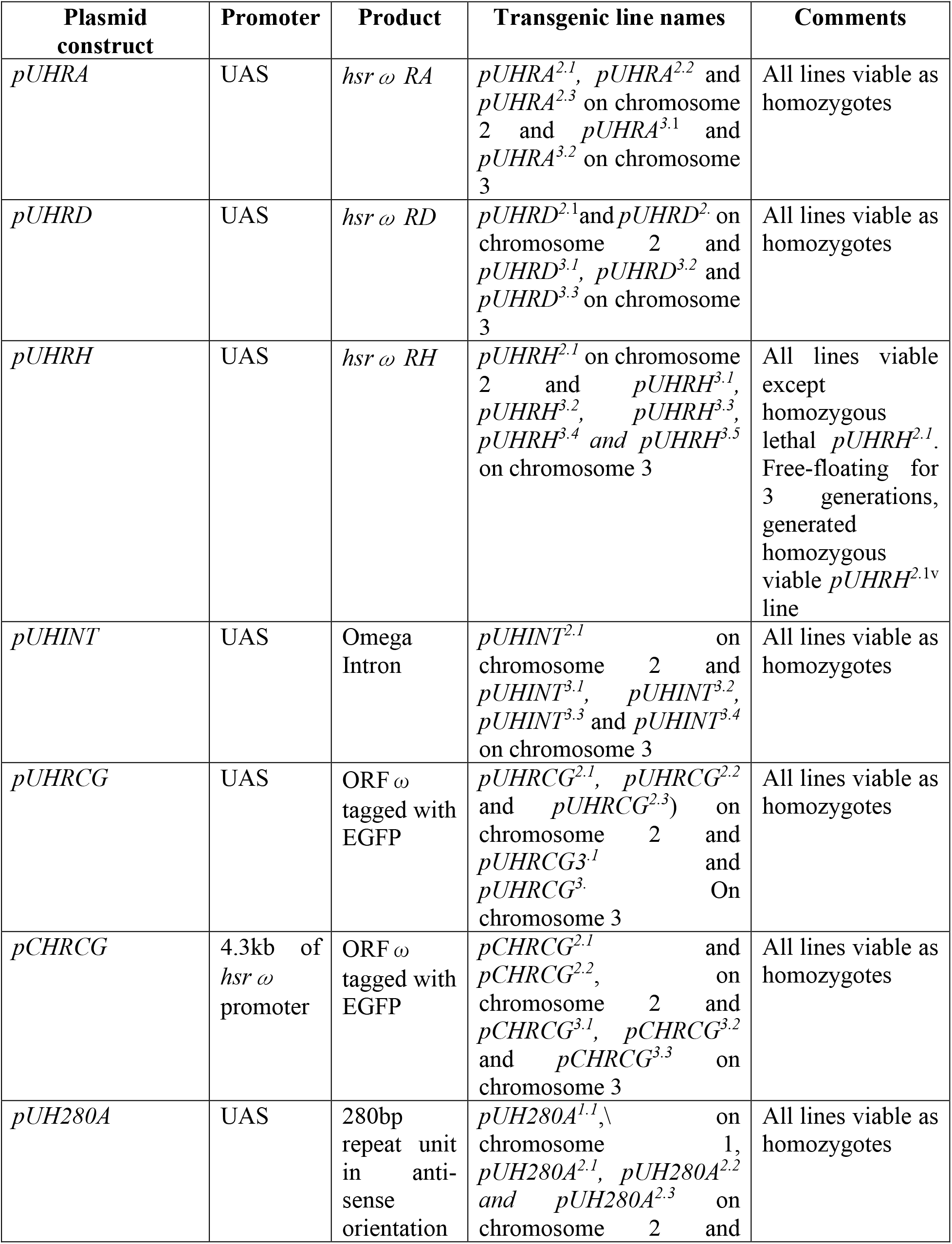
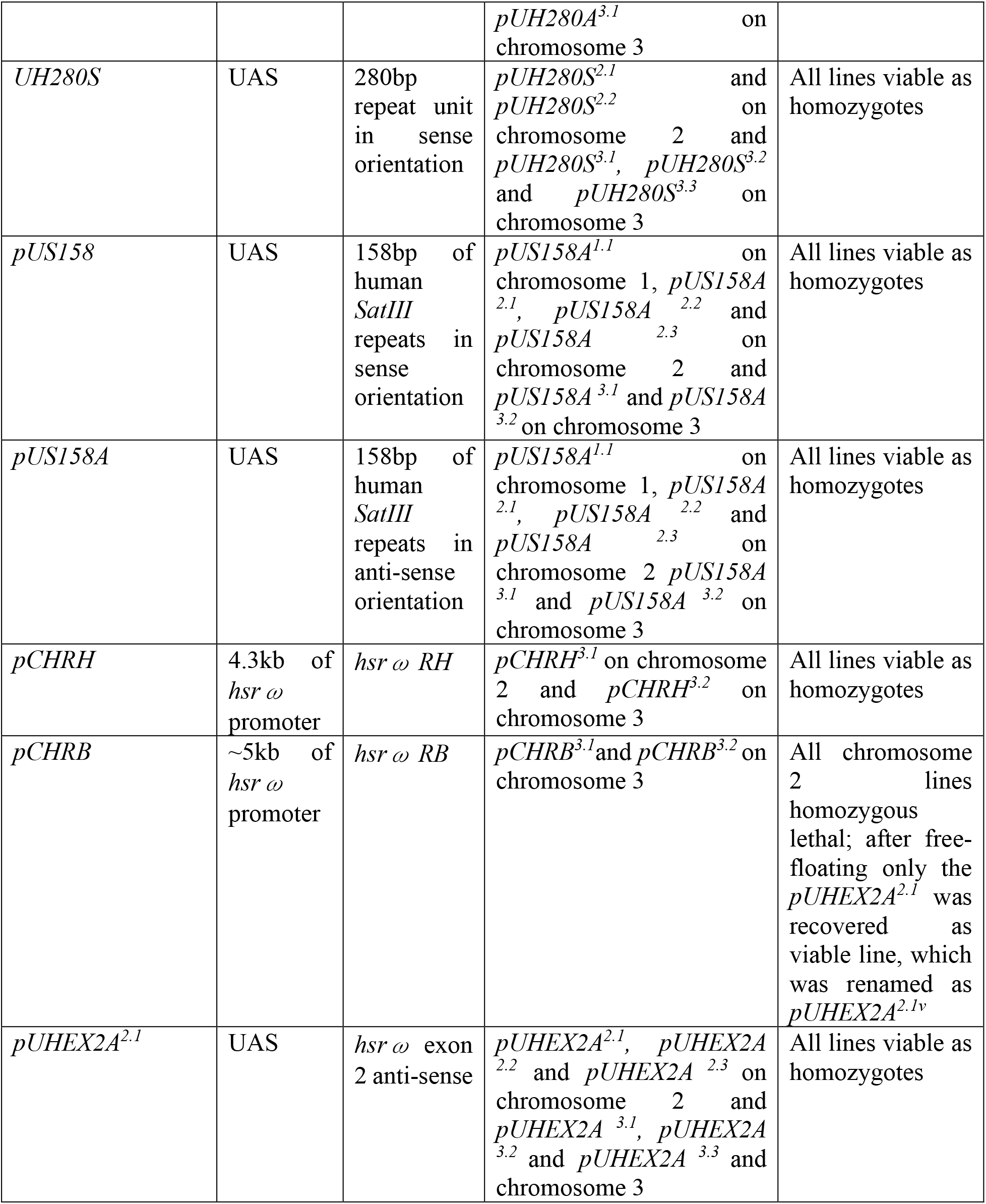
Summary of generated transgenic and gene-edited mutant *hsrω* alleles

**Table 2.**
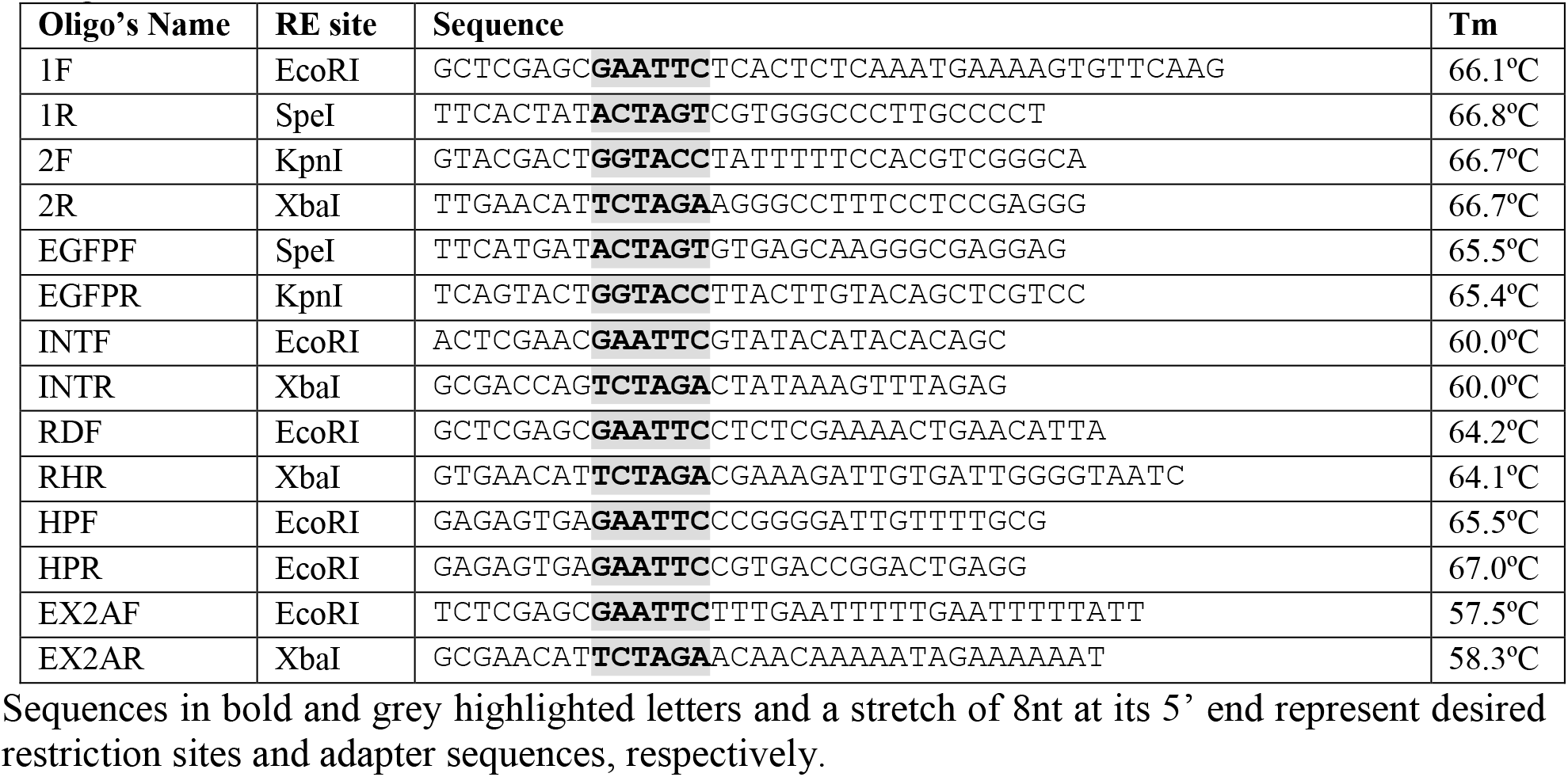
List of adapter primers used for amplification of inserts required for generation of transgenic constructs.

**Table 3.**
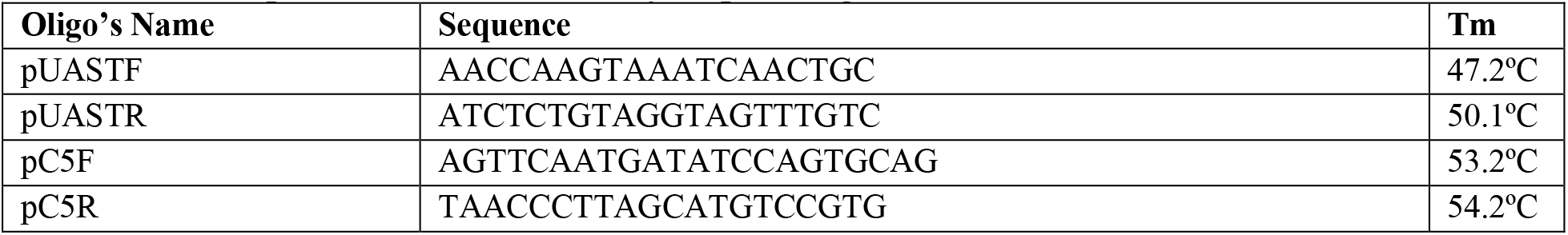
List of primers used for dideoxy sequencing of constructs.

### E. Transgenic constructs to generate CRISPR/Cas9 mediated ORFω knockout and complete *hsrω* gene knockout alleles

The CRISPR/Cas9 mediated genome editing was used to generate two deletion alleles of *hsrω* gene, one with the entire gene deleted, and the other with deletion of the ORFω region. The specific gRNAs (Table 4) for each, with the “NGG” sequences at their 3’ ends, were designed with the help of Benchling tool (www.benchling.com).

i. ***hsrω^KO^***: For complete deletion of the *hsrω* gene, the two gRNAs (Table 4) corresponded, respectively, to the 5’ region (gRNA-1) downstream of TSS1, and 3’ region (gRNA-2) upstream of the TTS4 (Fig. 2a). These were cloned in *pBFv-U6.2B* for generating a transgenic stock in which the transgene is inserted on chromosome II (*y^1^ v^1^, gRNA^1+2^/CyO*; +/+). The transgenic line carrying the gRNAs was crossed with flies expressing Cas9 in germline (*y*^1^ *v*^1^; *nos-Cas9*). The progeny F1 flies carrying the *gRNA* and *Cas9* transgenes were crossed with *w^1118^*; +/+; *TM3, Sb/TM6B* flies. The F2 progeny males were pair mated with *w^1118^*; +/+; *TM3, Sb/TM6B* females. After 2-3 days when larvae started appearing in the food vials, the parental males from each of the pair-mating vials were removed and their DNAs were isolated using the single fly DNA extraction method. Each of the F2 parental male DNA sample was screened for the desired deletion by PCR using the HKOF1 and HKOR1 primer pairs (Table 5). Only the DNA from which the *hsrω* gene is knocked out will generate a 674bp size amplicon while in wild type genomic DNA, the two primers anneal more than 18kb apart and thus fail to amplify the intervening DNA (Fig. 2b). The selected F3 flies were self-crossed to maintain the two *hsrω* knockout alleles (*hsrω^KO1^* and *hsrω^KO2^*). Absence of the gRNA or the Cas9 transgene in the deletion-positive flies was confirmed molecularly. Both these lines show complete embryonic/larval lethality when homozygous. They are maintained with a 3^rd^ chromosome balancer.
ii. ***ORFω^KO^:*** In order to generate ORFω knockout allele using the CRISPR/Cas9 technology, the 5’ gRNA corresponded to TSS2 region (gRNA-3, Table 4, Fig. 2a) while the 3’ gRNA (gRNA-4, Table 4, Fig. 2a) corresponded to exon 1 region downstream of the ORFω. These gRNAs were cloned in *pBFv-U6.2B* and used to generate transgenic stock (*y^1^ v^1^, gRNA^3+4^/CyO*; +/+). The transgenic progeny was crossed following the above scheme. PCR–based screening of ~300 F2 parental males using the primers RDUF and INTSR (Table 5) was carried out to identify males that carried the deletion of 268bp (Fig. 2b) and thus generated an amplicon of 543bp. Two independent transgenic lines (*ORFω^KO1^* and *ORFω^KO2^*) were obtained, both of which were homozygous viable but completely female sterile. A free floating of *ORFω^KO2^* allele carrying chromosome for three generations yielded progeny male and female flies that carried the *ORFω^KO2^* allele but were fertile. Presence of the *ORFω^KO2^* allele in this line was confirmed by PCR screening using the ORFω deletion detecting primers (RDUF and INTSR) and the allele was renamed as *ORFω^KO2f^*.

**Fig. 2.**
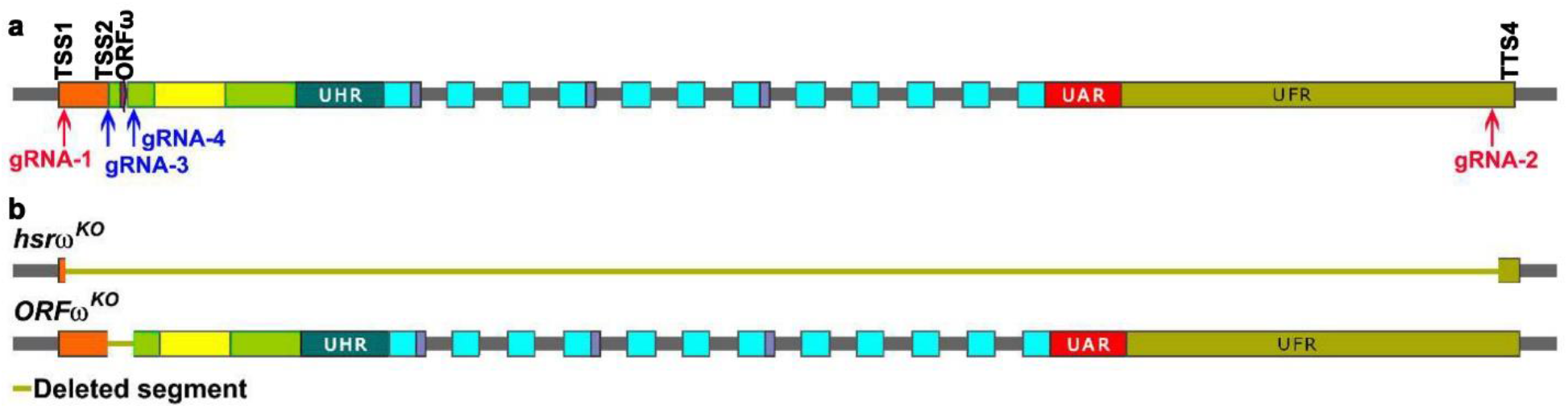
Generation of knockouts of *hsrω* using CRISPR/Cas9 system. **a**. Schematic of the *hsrω* gene with target locations of different gRNAs (gRNA-1 to gRNA-4) indicated in relation to the transcription start sites (TSS1 and TSS 2), ORFω and the last transcription termination site (TTS4). **b**. Maps of the *hsrω^KO^* and *ORFω^KO^* alleles carrying complete deletion of *hsrω* (from just after the TSS1 to just before the TTS4) or of the ORFω (from TSS2 to just after the ORFω in exon 1), respectively. The deleted regions are indicated by green line.

**Table 4.**
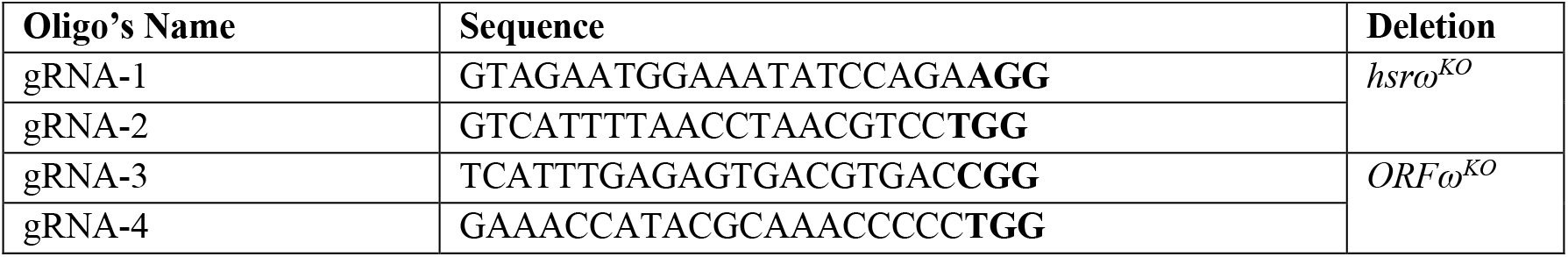
List of gRNAs used in generation of CRISPR/Cas9 knockouts (PAM site indicated in bold).

**Table 5.**
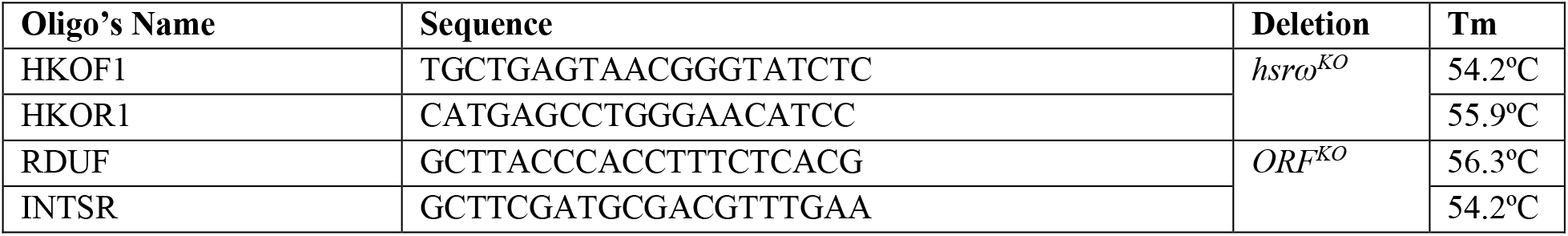
List of primers used for screening of CRISPR/Cas9 knockouts.

## Discussion

The various transgenic lines expressing *RD*, *RH* or *RB* transcripts under the *UAS* promoter can help in understanding functions of these transcripts through analysis of phenotypes induced by their targeted over-expression or by expressing them in *hsrω* null-background. The *hsrω^KO^* transgenic lines will be especially useful in this context. Likewise, the transgenic lines producing the *RH* or *RB* transcripts under the endogenous *hsrω* promoter will also be very useful when brought in *hsrω*-null background since in this case, unlike the *UAS*-promoter carrying transgene, each of these two transgenes are expected to express more or less in the normal developmentally regulated manner.

As noted above, the presently available *hsrω-RNAi* transgene primarily targets the 280bp repeat units in the nuclear transcripts and thus has little effect on the *hsrω* cytoplasmic transcripts (Mallik and Lakhotia, 2009). The *hsrω*-exon RNAi transgenic line generated in this study is expected to down-regulate all the transcripts of this gene since the exonic region is common to all the seven known transcripts; this RNAi line may have a greater effect on cytoplasmic *hsrω* transcript. The *hsrω* large nuclear repeat-carrying transcripts and the human *SatIII* transcripts are functional homologs (Jolly and Lakhotia, 2006). The two transgenic lines carrying the *SatIII* repeat unit in sense and anti-sense orientations, respectively, will thus be useful in comparing functions of the 280bp *hsrω* repeat unit and the 158bp *Sat III* repeat units and to see if these can complement each other.

We believe that the transgenic lines and the knockout alleles generated in this study will provide valuable resources and would indeed help researchers in a deeper understanding of the *hsrω* gene, which is one of the earliest known lncRNA gene (Lakhotia, 2011). Desiring users may write to the corresponding author for obtaining the required lines described here.

## ACKNOWLEDGEMENTS

This work has been supported by research grant from the Department of Biotechnology (Govt. of India) to SCL. RS was supported by a Research fellowship from the Council of Scientific and Industrial Research (Govt. of India). We thank the Fly Facility, Bangalore Life Science Cluster, Bangalore, for micro-injections in fly embryos to generate the transgenic lines.

## References

Giordano, E., R. Rendina, I. Peluso, and M. Furia 2002, Genetics 160: 637–648.

Jolly, C., and S.C. Lakhotia 2006, Nuclic Acids Research 34: 5508–5514.

Kang, S. G., E. Lee, S. Schaus, and E. Henderson 2001, Mo.l Med. 33: 174–178.

Lakhotia, S.C. 2011, J. Biosciences 36: 399–423.

Lakhotia, S.C. and A.S. Mukherjee 1970, Dros. Inf. Ser. 45: 108.

Mallik, M. and S. C. Lakhotia 2009, RNA Biol. 6: 464–478.

Prasanth, K., T. Rajendra, A. Lal and S.C. Lakhotia 2000, J. Cell Sci. 113: 3485–3497.

Sahu, R.K., E. Mutt and S.C. Lakhotia 2020, J. Genet. 99: 64.

Singh, A.K. and S.C. Lakhotia 2015, Chromosoma 124: 367–383.

Tapadia, M.G. and S.C. Lakhotia 1997, 5: 359–362.

Thummel, C.S., A.M. Boulet and H.D. Lipshitz 1988, Gene 74: 445–456.

